# Intercontinental movement of H5 2.3.4.4 Highly Pathogenic Avian Influenza A(H5N1) to the United States, 2021

**DOI:** 10.1101/2022.02.11.479922

**Authors:** Sarah N. Bevins, Susan A. Shriner, James C. Cumbee, Krista E. Dilione, Kelly E. Douglass, Jeremy W. Ellis, Mary Lea Killian, Mia K. Torchetti, Julianna B. Lenoch

## Abstract

Eurasian-origin highly pathogenic avian influenza A(H5N1) belonging to the Gs/GD lineage, clade 2.3.4.4b, was detected in two Atlantic states in wild waterfowl in the United States in January 2022. Bird banding data show widespread movement of waterfowl both within the Atlantic Flyway and between neighboring flyways and northern breeding grounds.

## Overview

Influenza A viruses (IAV) have a worldwide distribution and wild birds are the primary wild reservoir. Many wild ducks in particular are often repeatedly exposed to and infected with IAVs with little to no sign of clinical disease (1), although highly pathogenic (HP) forms of the virus can sometimes cause morbidity and mortality in wild birds (2). Highly pathogenic lineage viruses first identified in 1996 (A/goose/Guangdong/1/1996 [Gs/GD],) have repeatedly spilled over from poultry to wild birds and subsequent emergence of HP Gs/GD clade 2.3.4.4 viruses has led to circulation of these viruses in wild birds and high morbidity and mortality outbreaks in poultry on multiple continents (3).

One way to better understand IAV movement on the landscape or to identify routes of introduction of novel IAVs is through wild bird band-recovery data (4). These data have been collected as part of waterfowl management and conservation efforts in North America since the 1920s (5). Spatial locations of where birds are banded and later recovered are recorded and archived, providing data on wild bird movement. For waterfowl, recoveries primarily occur through banded individuals being reported subsequent to hunter harvest.

Wild bird samples are routinely collected by the USDA APHIS WS National Wildlife Disease Program (NWDP, USFWS Permit # MB124992 0) and screened for IAV in conjunction with the National Animal Health Laboratory Network and with the National Veterinary Services Laboratories (Ames, Iowa, USA) as part of a targeted IAV surveillance program in wild birds. These surveillance data combined with bird band-recovery movement data can help inform poultry producers, researchers, and government agencies on IAV occurrence on the landscape; and findings in wild birds can act as an early warning system for spillover risk to poultry.

For these findings, IAV viral RNA from wild bird samples was amplified as previously described (6). Following amplification, cDNA libraries were generated for MiSeq using the Nextera XT DNA Sample Preparation Kit (Illumina) and the 500 cycle MiSeq Reagent Kit v2 (Illumina) according to manufacturer instructions. De novo and directed assembly of genome sequences was carried out using IRMA v 0.6.7 (7) followed by visual verification in DNAStar SeqMan v14. For phylogenetic analysis, sequences were downloaded from GISAID and aligned in Geneious 11.1.5 using MAFFT and trees were generated using RAxML.

North American Bird Banding Program data (8) were queried to find all records from 1960-2021 for 11 dabbling duck species targeted by the NWDP wild bird surveillance. Targeted species included American black duck (*Anas rubripes*), American green-winged teal (*Anas crecca carolinensis*), American wigeon (*Mareca americana*), blue-winged teal (*Spatula discors*), cinnamon teal (*Spatula cyanoptera*), gadwall (*Mareca strepera*), mallard (*Anas platyrhynchos*), mottled duck (*Anas fulvigula*), northern pintail (*Anas acuta*), northern shoveler (*Spatula clypeata*), and wood duck (*Aix sponsa*). Records for these species were then limited to only include birds that were either banded or encountered in North Carolina or South Carolina and at least one other state or province.

## Results

As part of these routine surveillance efforts, in January 2022, HP Gs/GD lineage clade 2.3.4.4b H5N1 viruses were detected in multiple wild birds in North Carolina and South Carolina, USA (Figure 1). Genetic analyses demonstrate all virus segments are Eurasian origin (99.7-99.8% similar, Supplemental Information) and have high identity to December 2021 HP H5N1 findings in Newfoundland, Canada (9, Figure 1).

**Figure 1.**
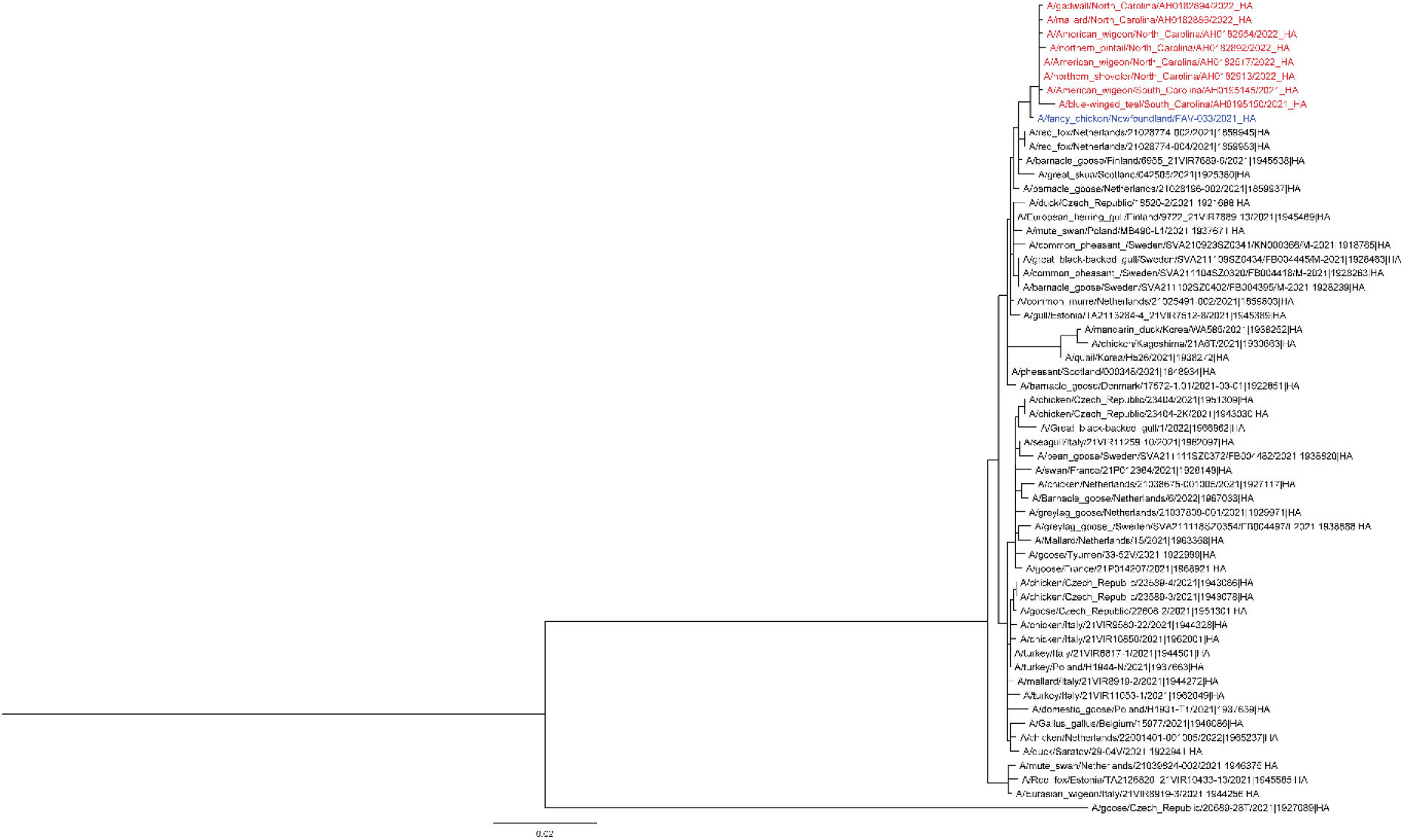
Maximum-likelihood phylogenic analysis of the hemagglutinin (HA) gene segment of the first set of wild bird isolates sequenced. Scale bars indicate average nucleotide substitutions per site. Outbreak viruses in red, closest virus detected in Newfoundland, Canada in blue.

The initial US detection was from a sample collected on December 30, 2021 from a wigeon in Colleton County, South Carolina (A/American_wigeon/South_Carolina/AH0195145/2021(H5N1), GISAID [https://www.gisaid.org] accession no. EPI_ISL_9869760). Immediately following this initial detection, there was an additional wild bird detection in South Carolina (A/blue-winged_teal/South_Carolina/AH0195150/2021(H5N1), GISAID accession no. EPI_ISL_9876777) and detections in neighboring North Carolina (Figure 1). Within six weeks there were 137 additional detections in wild birds, indicating high susceptibility to a novel virus and continued dispersal (Table). All birds were apparently healthy hunter harvested dabbling ducks. There was no detection of North American lineage IAV in any of these samples.

**Table:** The first detection of clade 2.3.4.4 HP IAV in the US was from a wild bird sampled on 30 December 2021. Within six weeks there were 137 additional detections in wild bids.

| **State** | **Wild bird species** | **Number of Clade 2.3.4.4.**<br>**HP IAV Detections** |
| --- | --- | --- |
| **South Carolina** | American wigeon | 7 |
|  | Blue-winged teal | 9 |
|  | Gadwall | 7 |
|  | Northern shoveler | 1 |
| **North Carolina** | American green-winged teal | 9 |
|  | Northern shoveler | 3 |
|  | American wigeon | 51 |
|  | Gadwall | 15 |
|  | Mallard | 4 |
|  | Northern pintail | 3 |
|  | Wood duck | 1 |
| **Virginia** | American green-winged teal | 2 |
|  | Mallard | 1 |
| **Florida** | Blue-winged teal | 2 |
| **Delaware** | American wigeon | 1 |
|  | Northern shoveler | 2 |
| **New Hampshire** | Mallard | 20 |
| **Total Detections** |  | 138 |

Analysis of North American Bird Banding Program data show broadscale movement of waterfowl throughout North America. Across 11 species of dabbling ducks targeted in surveillance sampling that were banded or encountered in North Carolina or South Carolina and subsequently or previously banded or encountered in another state or province, 64.7% of bird movements were within the Atlantic Flyway, 33.6% of analyzed species were encountered in both the Atlantic and the Mississippi Flyways, and 1.7% were encountered in both the Atlantic and Central Flyways (Figure 2).

**Figure 2.**
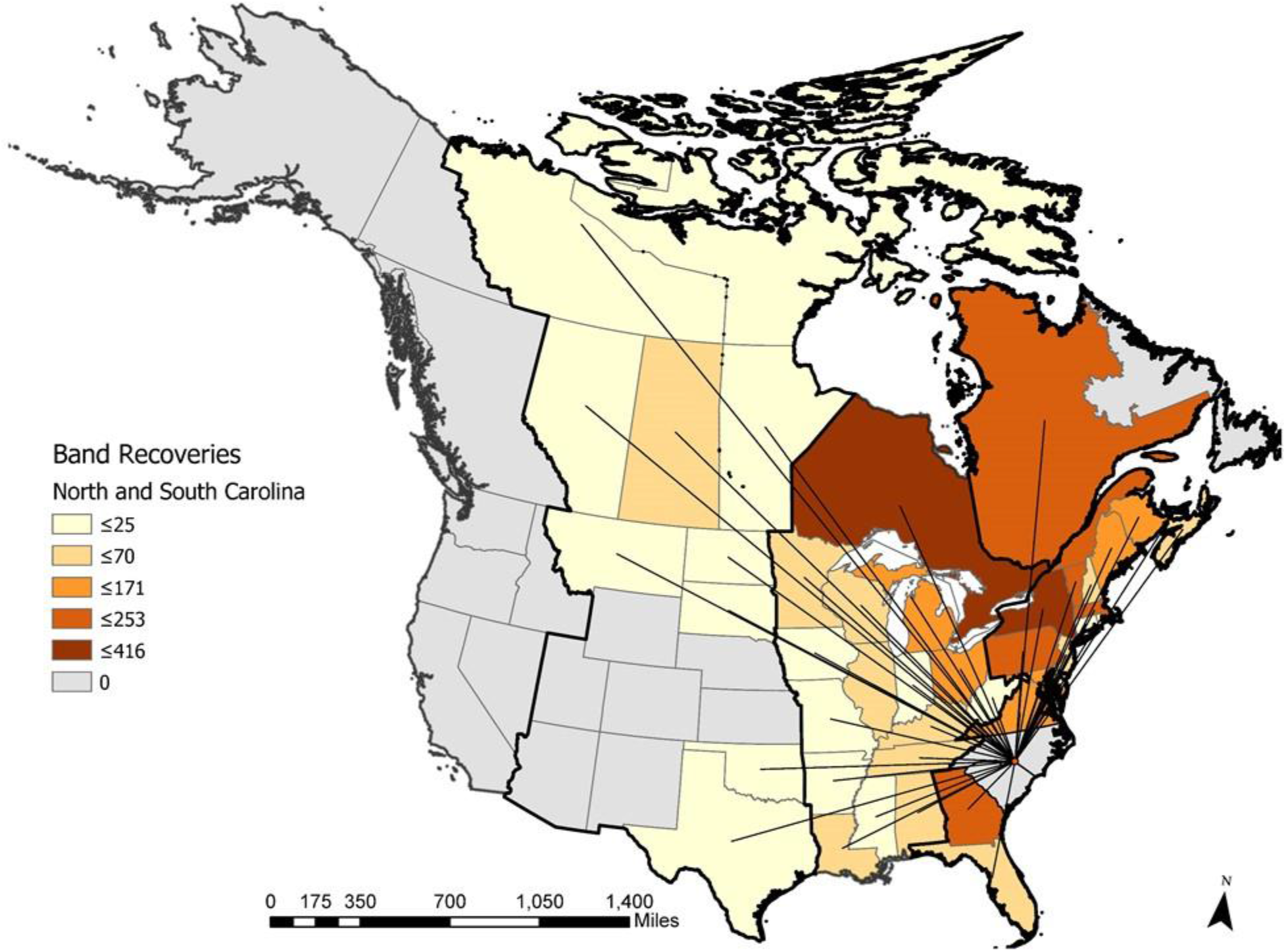
Map of dabbling duck movements to and from North Carolina and South Carolina to other states or provinces. Data are based on North American Bird Banding Program data collected between 1960-2021. Color intensities represent the number of movements detected between a given state or province and North Carolina or South Carolina. Lines are positioned at the centroid of a given state or province. Bold border lines represent the administrative migratory bird flyways (from West to East: Pacific Flyway, Central Flyway, Mississippi Flyway, and Atlantic Flyway).

## Conclusions

While there had been intense focus on intercontinental movement of HP IAV from Asia to the North American Pacific Flyway (10), viral movement via the trans-Atlantic pathway has been less clear. Data reported here, in combination with the recent HP findings in Newfoundland, Canada (9), point to this being the first documented introduction of a Eurasian origin HP IAV into wild birds via the Atlantic Flyway of the US. The potential introduction pathway likely includes wild bird migratory routes from Northern Europe that overlap Arctic regions of North America, and then dispersal farther south into Canada and the US (11,12).

Band recovery data reveal a majority of dabbling ducks banded in the Atlantic Flyway are also recovered in the Atlantic Flyway, reinforcing the predominance of within flyway movement (13); however, data also reveal routine movement to other flyways, providing a potential mechanism of wider spread dispersal of the virus in North American waterfowl. In addition, the sequence data indicate that these viruses cluster closely with viruses found in Western Europe during the spring of 2021 (Figure 1, Supplementary Information), offering the possibility that these viruses are already in multiple locations in North America. Additional detections in wild birds suggest these 2.3.4.4b H5 viruses continue to be transmitted (Table S1, Supplemental Information) and further dispersal may be seen once waterfowl begin migration to summer breeding areas.

Many previous HP findings in wild birds have been tied to repeated spillover of the viruses from domestic birds, which is where mutations to high pathogenicity primarily occur; however, in some cases, Gs/Gd lineage viruses now appear to be maintained in wild bird populations (14). This potential adaption of HP IAV to wild birds highlights the importance of continued wild bird surveillance in the future. In addition, these findings demonstrate that targeted IAV surveillance in wild bird populations can detect newly introduced or emergent IAV viruses prior to spillover to domestic poultry. Advanced warnings from wild bird surveillance allow poultry producers to consider altering biosecurity in the face of increased IAV risk and also help inform zoonotic disease potential (15).

## Acknowledgements

We thank Wildlife Services employees and collaborators at state wildlife agencies for contributing wildlife sampling expertise to this large-scale effort. We also thank the staff at state and federal agency laboratories and at the Canadian Food Inspection Agency that contributed to this study. Funding provided by U.S. Department of Agriculture, Animal Plant Health Inspection Service.

## Disclaimer

Any use of trade, firm, or product names is for descriptive purposes only and does not imply endorsement by the US Government.

## Biographical Sketch

Dr. Bevins and Dr. Shriner are Research Scientists at the USDA APHIS WS National Wildlife Research Center who work on pathogen emergence from wild animals.

