## Supplementary Information for "Intercontinental movement of H5 2.3.4.4 Highly Pathogenic Avian Influenza A(H5N1) to the United States, 2021"

### Supplemental Information

#### Supplemental Figure Legend

Fig S1. Maximum-likelihood phylogenic analysis of the PB1 gene segment. Scale bars indicate average nucleotide substitutions per site. Outbreak viruses in red, closest detection in Newfoundland, Canada in blue.

Fig S2. Maximum-likelihood phylogenic analysis of the PB2 gene segment. Scale bars indicate average nucleotide substitutions per site. Outbreak viruses in red, closest detection in Newfoundland, Canada in blue.

Fig S3. Maximum-likelihood phylogenic analysis of the MP gene segment. Scale bars indicate average nucleotide substitutions per site. Outbreak viruses in red, closest detection in Newfoundland, Canada in blue.

Fig S4. Maximum-likelihood phylogenic analysis of the NA gene segment. Scale bars indicate average nucleotide substitutions per site. Outbreak viruses in red, NA gene segment detected in Newfoundland, Canada in blue.

Fig S5. Maximum-likelihood phylogenic analysis of the NP gene segment. Scale bars indicate average nucleotide substitutions per site. Outbreak viruses in red, closest detection in Newfoundland, Canada in blue.

Fig S6. Maximum-likelihood phylogenic analysis of the NS gene segment. Scale bars indicate average nucleotide substitutions per site. Outbreak viruses in red, closest detection in Newfoundland, Canada in blue.

Fig S7. Maximum-likelihood phylogenic analysis of the PA gene segment. Scale bars indicate average nucleotide substitutions per site. Outbreak viruses in red, closest detection in Newfoundland, Canada in blue.

#### Figure S1

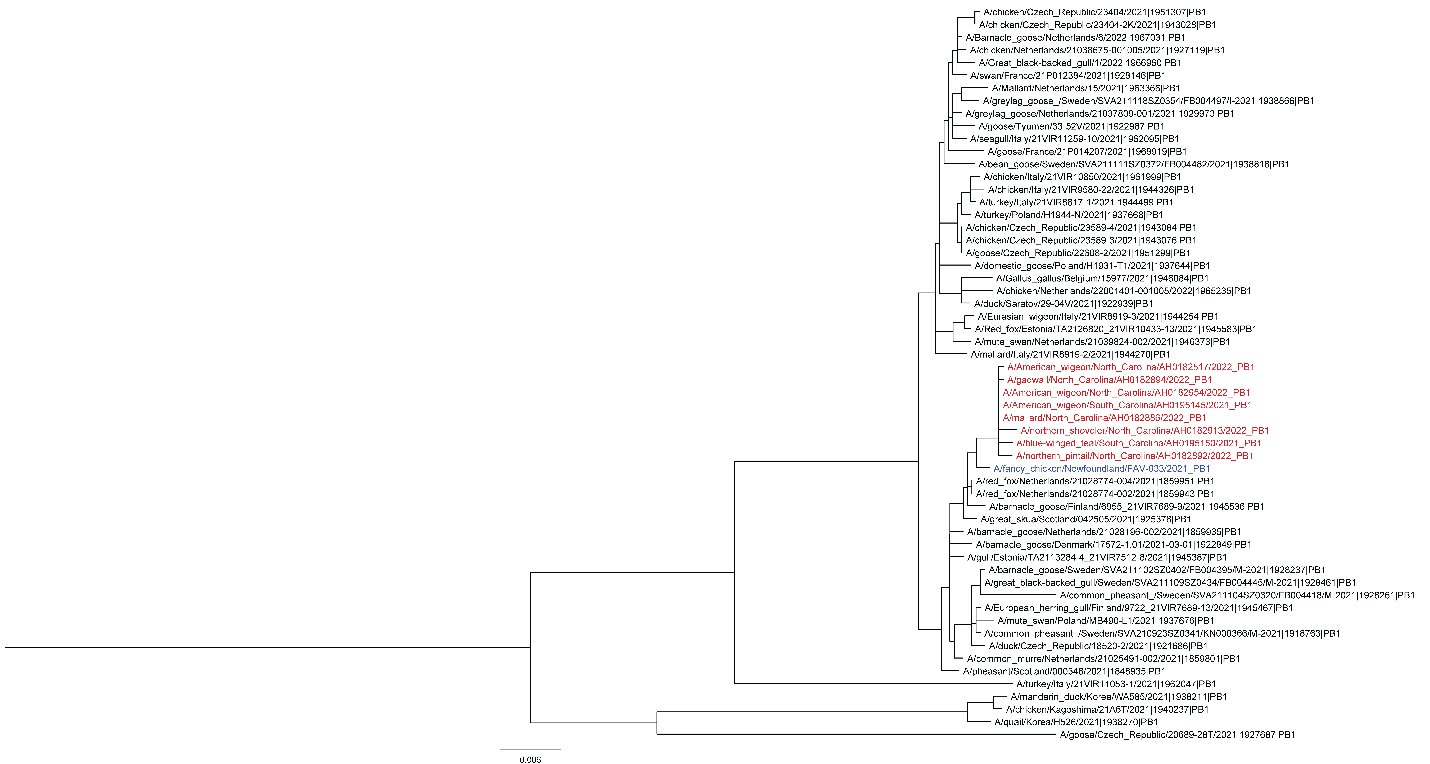

#### Figure S2

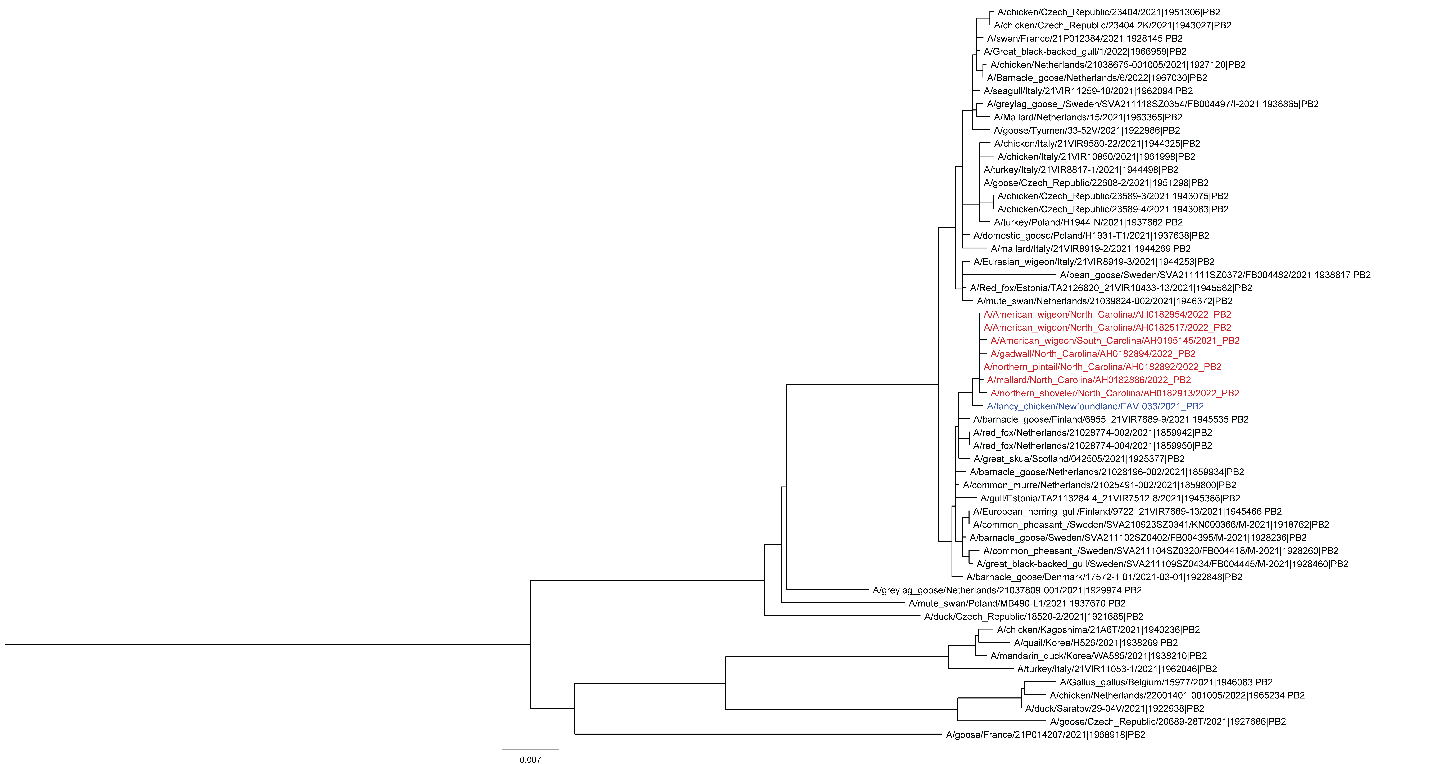

#### Figure S3

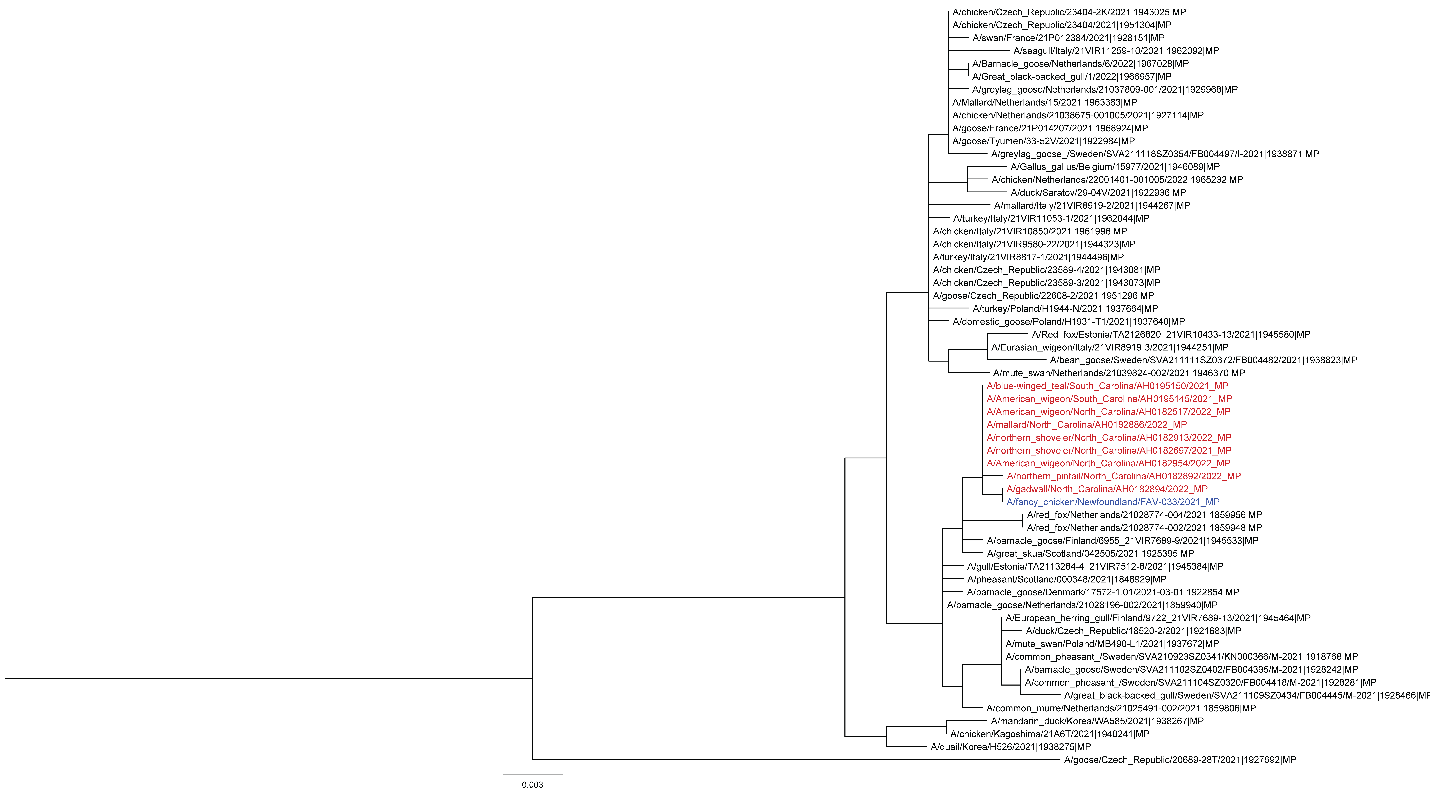

#### Figure S4

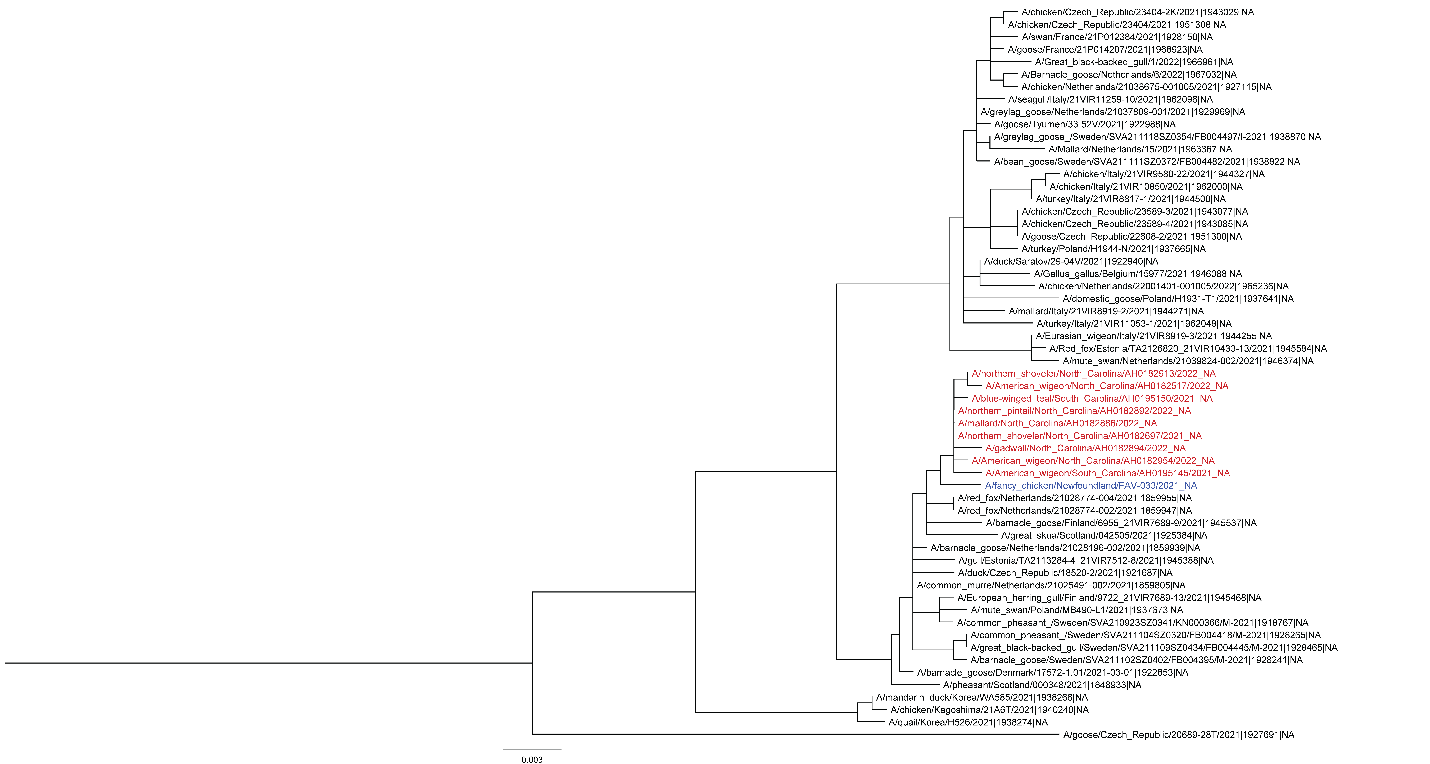

#### Figure S5

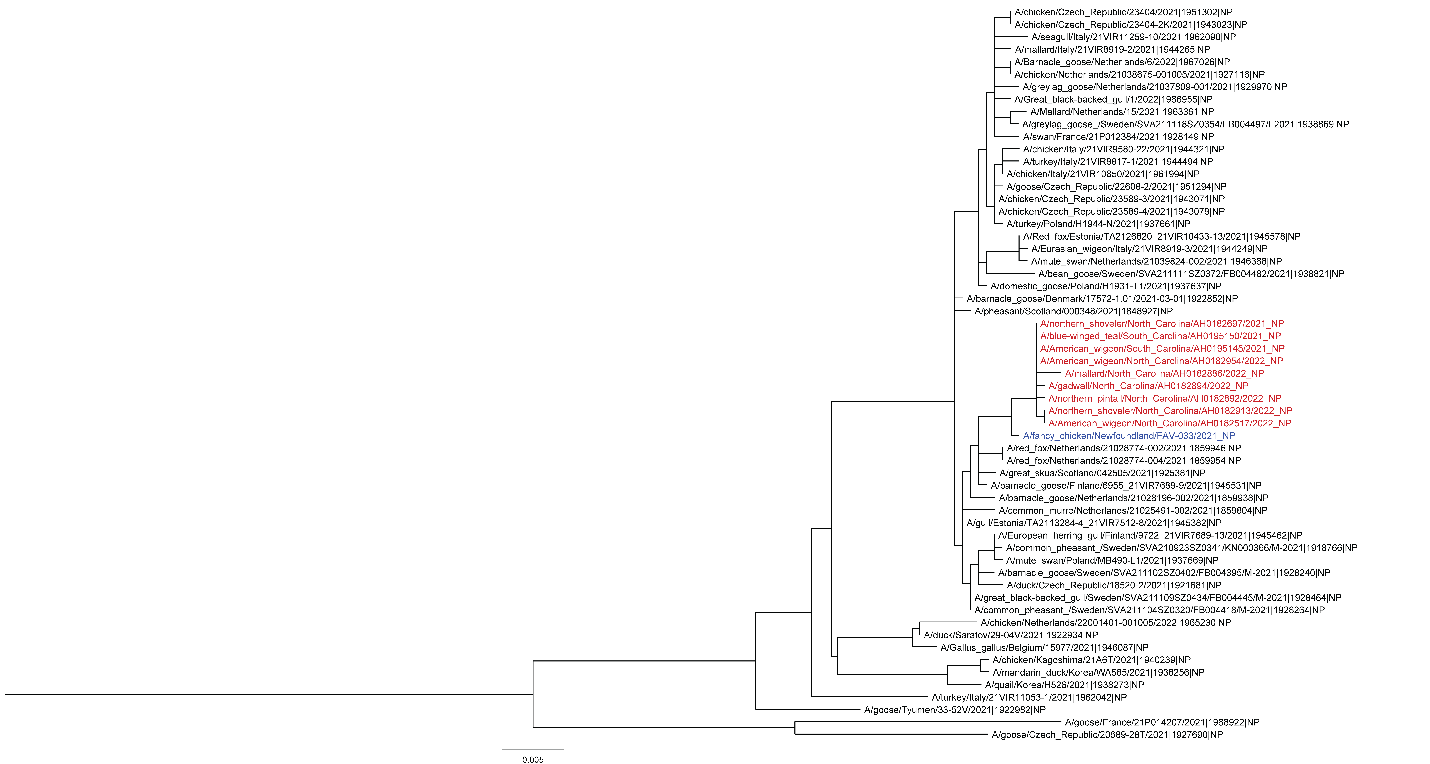

#### Figure S6

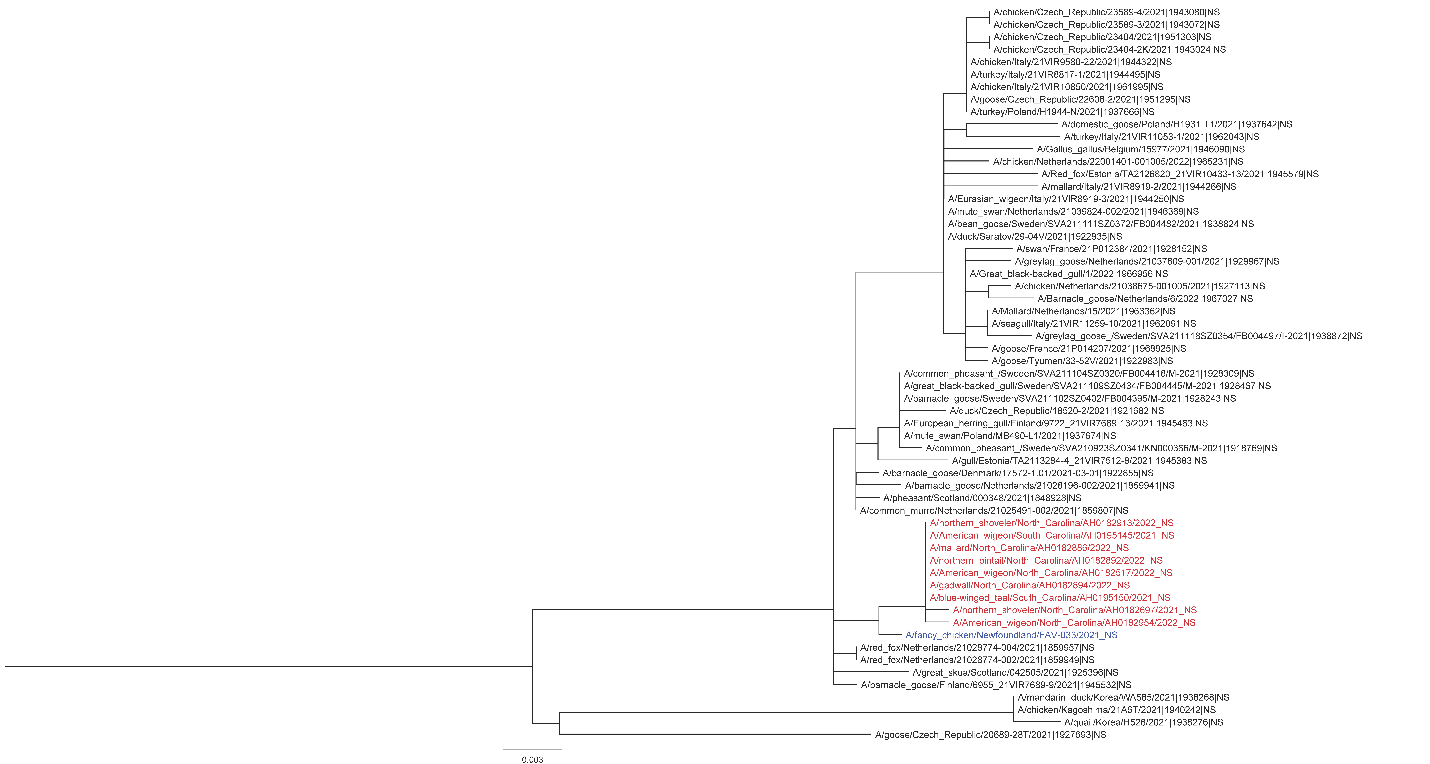

#### Figure S7

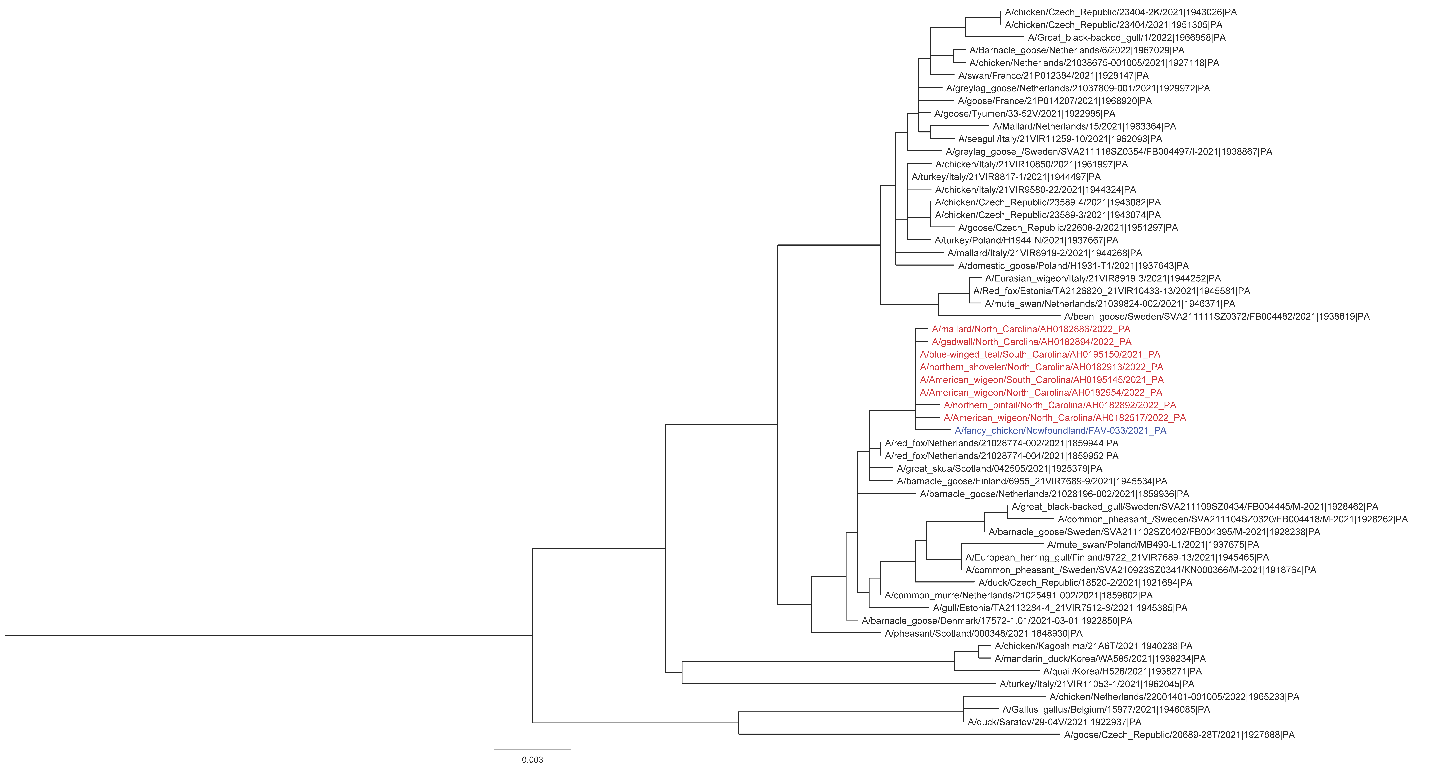
